# Conservation status, threats, and information needs of small mammals in Alaska

**DOI:** 10.1101/2020.10.11.330555

**Authors:** A. Droghini, K. S. Christie, R. R. Kelty, P. Schuette, T. A. Gotthardt

**Author notes:** **Corresponding author:** A. Droghini.

## Abstract

Despite their diversity and ecological importance, small mammals are under-represented in conservation research relative to other mammals. We evaluated the conservation status of 36 small mammal species in Alaska, U.S.A. using a ranking system that we previously developed, the Alaska Species Ranking System (ASRS). We compared results from the ASRS with NatureServe’s subnational rankings. Finally, we surveyed taxonomic experts to identify recommended conservation actions and research priorities for 5 species of high conservation concern. In general, the ASRS and NatureServe agreed on the rankings of species in the highest and lowest risk categories. Species of highest conservation concern were taxa endemic to the state, including 2 island-endemic shrews, and taxa from the orders Chiroptera and Eulipotyphla. Because the ASRS includes information needs in its assessment, 15 of the 20 species considered lowest concern by NatureServe were considered intermediate concern by the ASRS. In the ASRS, most species (*n* = 24) were assessed to have low biological vulnerabilities, but high information needs. Population size and trends were unknown for all species; distributional limits and understanding of population dynamics were incomplete for all species except 4. Disease and climate change effects on habitat were perceived as important threats, but affected only 8 species. Taxonomic experts identified addressing data deficiencies and protecting habitat as important conservation actions; they identified monitoring population trends, modeling habitat, and researching species’ genetic diversity and adaptive capacity as high priorities. Conservation assessments that require accurate and current data on population trends or threats may lead to bias against data deficient groups such as small mammals. Our findings demonstrate the importance of accounting for data deficiencies in conservation status ranks to avoid conflation of sparse information with low conservation concern.

## INTRODUCTION

Conservation practitioners and natural resource managers are often tasked with prioritizing effort and funding for species based on extirpation risk or vulnerability to threats. To aid with prioritization, practitioners often use ranking systems, such as those developed by the International Union for the Conservation of Nature (IUCN) or NatureServe. Conservation ranking systems assign a status to a species by evaluating and scoring that species across a set of criteria (IUCN 2012; Master et al. 2012). Scoring criteria requires data that are accurate, current, and available. When data are scarce or absent, species may receive a special status (e.g., data-deficient for the IUCN or unrankable for NatureServe). In less extreme cases, assessors can score some questions as unknown or select a range of answers to express uncertainty (IUCN 2012; Master et al. 2012). Designations of uncertainty allow assessors to assign conservation status to species that lack the data necessary to reliably score a subset of criteria; however, ranking systems can assign low-risk status to species with unknown population trends or threats if designations of uncertainty have no influence on the rank calculation. Data deficient species present challenges to conservation practitioners because the funding necessary to address data gaps can be difficult to justify for species with low-risk status as compared with species ranked as at-risk because of more complete or accurate data (Jetz and Freckleton 2015).

Small mammals (Chiroptera, Eulipotyphla, Rodentia, Lagomorpha) compose over 75% of the Earth’s extant mammalian diversity and function as primary consumers, insectivores, vectors of disease, and focal prey species (Ceballos & Brown 1995; Entwistle & Stephenson 2000). Despite their diversity and ecological importance, limited knowledge of population sizes, population and distribution trends, and threats often precludes assessment of their conservation status using traditional ranking systems. Under the IUCN ranking system, 16% of Rodentia species are listed as data-deficient, compared with 7% of Carnivora species and 4% of Cetartiodactyla species; however, a model developed by Jetz and Freckleton (2015) predicted more than half of the data deficient Rodentia species to be threatened. Small mammals are under-represented in conservation literature and receive less funding than other mammal groups (Entwistle & Stephenson 2000), likely resulting in the high proportion of small mammal species considered data deficient. Despite a lack of research attention and funding, most mammal species that have gone extinct in the past 500 years have been small mammals (Ceballos and Brown 1995). Thus, the conservation attention afforded to small mammals is indicative of neither their information needs nor their resilience.

The conservation status of species at the global scale does not reflect conservation threats and vulnerabilities at local scales (Breininger et al. 1998; Hartley and Kunin 2003). Thus, many jurisdictions develop their own ranking systems (Millsap et al. 1990; Breininger et al. 1998; Gotthardt et al. 2012). In 2007, the Alaska Department of Fish and Game Threatened, Endangered, and Diversity Program identified the need for a state-specific ranking system to evaluate the conservation status of tetrapods in Alaska. It partnered with the state’s Natural Heritage Program, the Alaska Center for Conservation Science (ACCS), to create the Alaska Species Ranking System (ASRS). The ASRS was modeled after a ranking system developed for the state of Florida (Millsap et al. 1990) and was modified to be relevant to Alaska’s ecological conditions and user needs.

Alaska has a unique geography and glacial history that has resulted in the evolutionary divergence of many taxonomic groups, including small mammal taxa that do not occur anywhere else in the United States (Cook et al. 2001; Lanier et al. 2015). Unlike other states in the U.S., threats from human development are low. However, in recent years, there has been concern about the effects of climate change to habitats, disturbance regimes, and species’ assemblages (Tape et al. 2006; Chapin et al. 2010; Tape et al. 2016).

In this paper, we use assessments from 2 conservation ranking systems, the ASRS and NatureServe, to summarize the status, threats, and data deficiencies of small mammal species in Alaska. We also elicited expert opinion to identify conservation actions and research priorities. By synthesizing results from 3 sources, we provide a comprehensive assessment of the conservation status of nearly all small mammal species found in Alaska.

## METHODS

From 2017 to 2020, we assessed the conservation status of 36 small mammal species in Alaska using the ASRS and the NatureServe Conservation Status Assessment. ACCS is part of the NatureServe network of Natural Heritage Programs and Conservation Data Centers (https://www.natureserve.org/); thus, ACCS is responsible for maintaining sub-national conservation status ranks for the state of Alaska.

### The Alaska Species Ranking System

The ASRS was developed in 2007 by ACCS and the Alaska Department of Fish & Game. Gotthardt et al. (2012) describe the ASRS in detail; we provide a brief summary here. The ASRS contains 13 multiple-choice questions classified into 3 themes: Trends, Biological Vulnerability, and Action Needs. The Trends theme comprises 2 questions evaluating change in population and distribution. The Biological Vulnerability theme characterizes the ecological and biological traits that correlate with extirpation risk for a species (Gotthardt et al. 2012). The Action Needs theme measures the strength of management and conservation actions. In this context, conservation actions evaluate knowledge gained from inventory, monitoring, and research efforts. In combination, the 3 themes effectively assess the risk of regional extirpation of non-endemic species and extinction of endemic species (collectively referred to as “conservation concern”).

Conservation concern increases with numeric value in the ASRS and scores can be positive or negative. The 2 questions in Trends are evaluated on a 5-point scale ranging from −10 to 10. A high Trends score indicates currently declining populations or shrinking distributions. Biological Vulnerability comprises 7 questions, which are evaluated on 3-, 4-, or 5-point scales. Three-point scales range from −5 to 5; the others range from −10 to 10. A high score for this theme indicates that the species has several traits (e.g., small population size, restricted range, slow life history, or high ecological specialization) that make it more vulnerable to extirpation. Finally, Action Needs comprises 4 questions that are each evaluated on a 3-point scale, which indicate low (−10), moderate (2), or high needs (10). A high Action Needs score denotes an absence of management and conservation actions, resulting in large information needs. Scores within themes are summed to create a theme score. Each theme score is then categorized to create a final, categorical rank.

The ASRS categorizes each theme score as high, unknown, or low; score thresholds for each category are presented in Gotthardt et al. (2012). Categorization results in 9 numerical ranks ranging from I to IX. Numerical ranks are further grouped into one of 4 color categories: red (highest conservation concern, numerical ranks I-II), orange (ranks III-V), yellow (ranks VII and VIII), and blue (ranks VI and IX). Red, Orange, and Yellow indicate high, unknown, or low Trends scores, respectively. Species in these categories also scored high for one or both of the remaining themes, Biological Vulnerability and Action Needs. One exception is the rank Orange III, which indicates a high Trends score but low Biological Vulnerability and Action Needs. Finally, a rank of Blue indicates low Biological Vulnerability and Action Needs, and either an unknown or low Trends score.

The assessment process for the ASRS begins with a trained assessor conducting a systematic review, preparing a species’ account, and conducting an initial assessment to obtain preliminary scores (Fig. 1). The assessor searches primarily for information relevant to Alaskan populations but expands the search to include other populations when information on Alaskan populations is insufficient to assign a score. A second assessor reviews the account and conducts an assessment without consulting the scores of the first assessor. If assessors disagree on a score, they discuss the question and consult additional sources. The assessment is then sent to a taxonomic expert for external review (Fig. 1). Once the review is complete, assessors update scores following the expert’s recommendation and finalize the assessment. Species’ accounts, along with related conservation resources, are published online where they are publicly available (Fig. 1).

**Figure 1.**
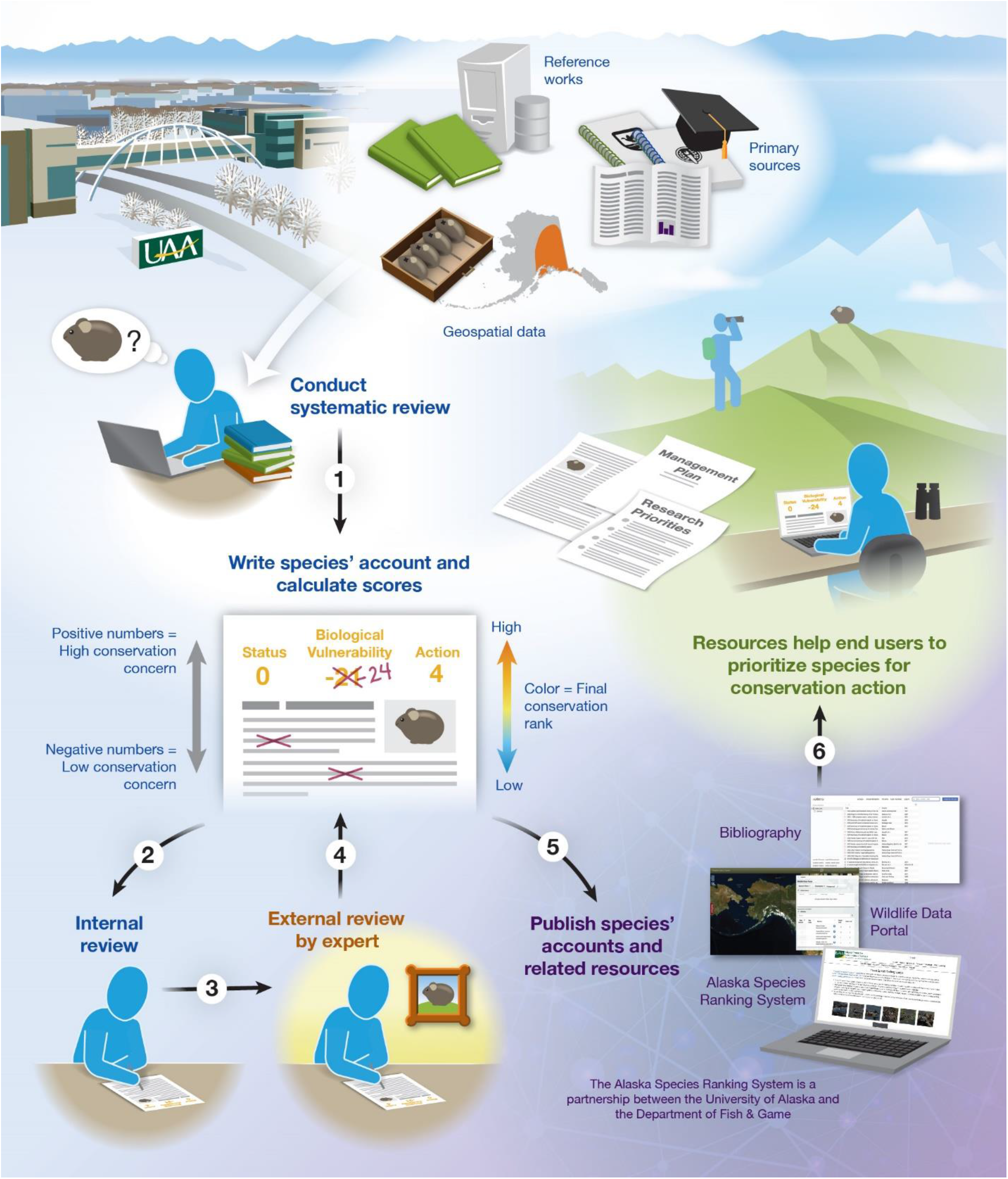
The Alaska Species Ranking System is a conservation ranking system for tetrapods in Alaska. It uses a multi-step process to ensure that assessments are objective, transparent, and standardized across taxa.

### NatureServe Conservation Status Assessments

NatureServe sub-national ranks (S ranks) are calculated by assessing a species’ Rarity, Population Trends, and Threats (Master et al. 2012). Criteria for Rarity include population size, range extent, area of occupancy, and the ecological integrity of known habitat. Population Trends consider short- and long-term trends. Finally, Threats are assessed by evaluating their scope (percent of population affected), severity (within the scope, percent of population reduction), and timing. An overall threat impact score is calculated using severity and scope scores across all the threats that were identified (Master et al. 2012).

S ranks range from 1 to 5. A rank of S1 indicates that the species is critically imperiled in that subnational jurisdiction (typically a state or province). A rank of S5 indicates that the species is secure in that subnational jurisdiction. A range rank (e.g., S3S4) indicates that the status of the species is uncertain within the bounds of the two values. Because the ASRS and NatureServe ranking systems use similar criteria, we used information from the ASRS species’ accounts to update the S ranks of the 36 small mammal species. Scientific literature and expert opinion informed the Threats assessments, which are not included in the ASRS (see next section).

### Identifying Conservation Actions and Research Priorities Using Expert Opinion

In 2019, we surveyed taxonomic experts to obtain their judgment on the most important conservation actions, research priorities, and threats facing 5 species: *Glaucomys sabrinus*, *Marmota broweri*, *Myotis lucifugus*, *Ochotona collaris*, and *Synaptomys borealis*. Each of the selected species are of high conservation concern (ADF&G 2015; this paper) and have been the topic of dedicated research projects in the state. For each species, we identified 3 to 7 experts. We defined an expert as a scientist who was directly involved in a research project investigating the species in Alaska.

Using an online survey tool, we asked experts a series of 7 questions (Table 1; Droghini et al. 2020). Answers to the first 5 survey questions informed the evaluation of threats for the 5 selected species and species with similar ecologies. The last 2 questions asked experts to identify conservation actions to mitigate threats and identify the most critical research needs if they were responsible for allocating a large sum of money (US $10 million) to research activities (Table 1).

**Table 1.**
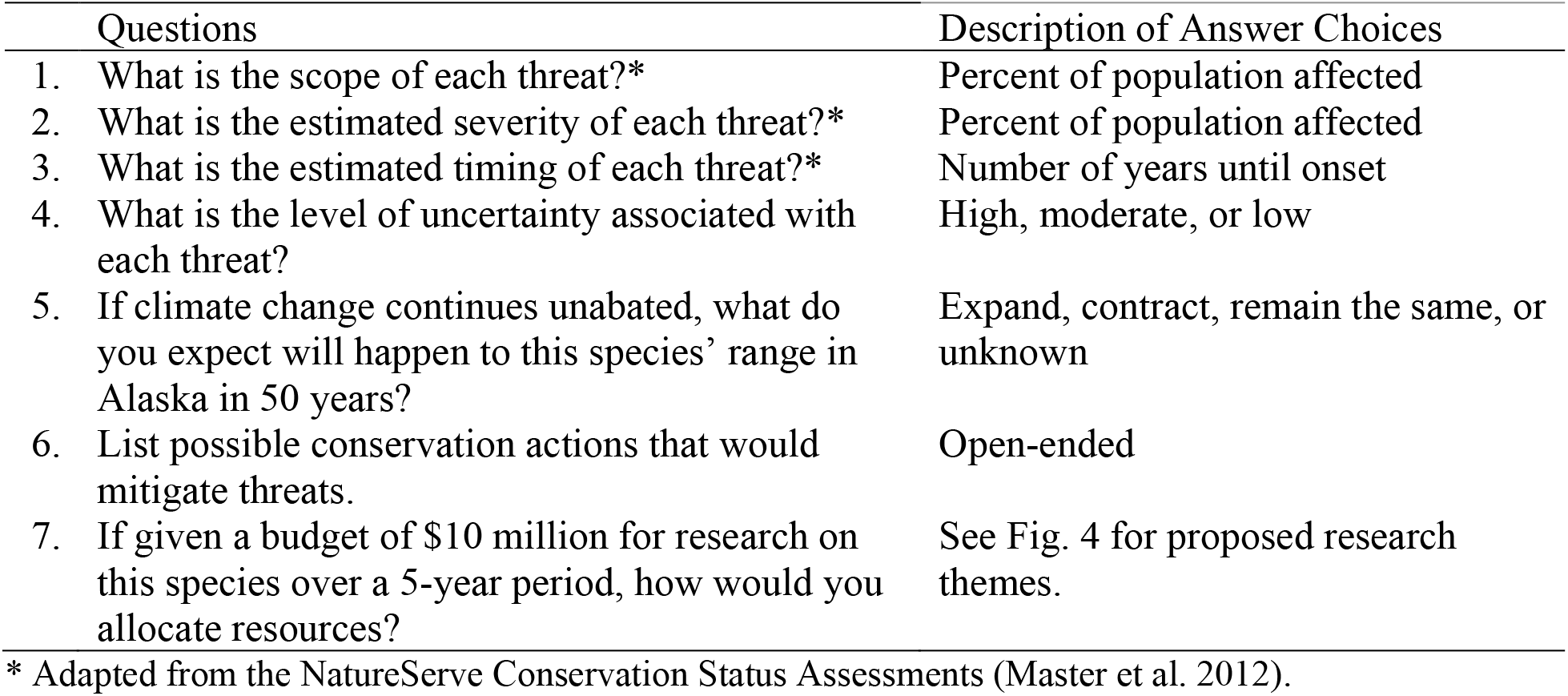
Questions posed to taxonomic experts to assess threats, conservation actions, and research priorities for 5 species of high conservation concern in the state of Alaska.

| Questions | Description of Answer Choices |
| --- | --- |
| 1. What is the scope of each threat?* | Percent of population affected |
| 2. What is the estimated severity of each threat?* | Percent of population affected |
| 3. What is the estimated timing of each threat?* | Number of years until onset |
| 4. What is the level of uncertainty associated with each threat? | High, moderate, or low |
| 5. If climate change continues unabated, what do you expect will happen to this species' range in Alaska in 50 years? | Expand, contract, remain the same, or unknown |
| 6. List possible conservation actions that would mitigate threats. | Open-ended |
| 7. If given a budget of \$10 million for research on this species over a 5-year period, how would you allocate resources? | See Fig. 4 for proposed research themes. |

\* Adapted from the NatureServe Conservation Status Assessments (Master et al. 2012).

## RESULTS

We evaluated the conservation status of 36 small mammal species across 4 orders and 7 families. Assessments from both the ASRS and NatureServe ranking systems are available online (Droghini et al. 2020). Relative to Alaska’s species diversity, we assessed 19 out of 23 Rodentia species, all Lagomorpha species (*n* = 3), all Eulipotyphla species (*n* = 9), and 5 out of 7 Chiroptera species. Four species are endemic to Alaska: *Lepus othus, Marmota broweri*, *Sorex jacksoni*, and *S. pribilofensis*. *Sorex jacksoni* and *Sorex pribilofensis* are endemic to the island of Saint Lawrence and Saint Paul, respectively; *L. othus* and *M. broweri* are more widespread. The taxonomy of *L. othus* and *S. jacksoni*, as well as *Myotis evotis*, are the subject of ongoing research (Waltari and Cook 2005; Hope et al. 2012; Cason et al. 2016; Lausen et al. 2019). Our focus was to assess the conservation status of species over which the state of Alaska has significant stewardship. The species we did not assess either have very restricted ranges in Alaska (e.g. *Zapus princeps*) or are not typically considered small mammals (e.g. *Castor canadensis*).

### Overall Status Ranks by ASRS and NatureServe

The highest rank obtained by a small mammal species in the ASRS was Orange IV. Orange IV indicates unknown Trends, high Biological Vulnerability, and high Action Needs (Gotthardt et al. 2012) and is the highest assignable rank for species that have unknown population and distribution trends. We assigned 7 species the rank of Orange IV: 3 species endemic to Alaska (*Marmota broweri, Sorex jacksoni*, and *S. pribilofensis*) and 4 Chiroptera species largely restricted to Southeast Alaska (*Lasionycteris noctivagans*, *Myotis californicus*, *M. evotis*, and *M. volans*). Under the NatureServe system, these 7 species received similar ranks of high concern relative to other species (Droghini et al. 2020). Specifically, *Marmota broweri, Sorex jacksoni*, and *S. pribilofensis* received a rank of S3 (vulnerable) when assessed using the NatureServe methodology; this rank was the highest rank obtained by the species we assessed.

Most species (*n* = 24) in the ASRS, including most Rodentia species (*n* = 14) and Eulipotyphla species (*n* = 7), ranked as Orange V, defined as unknown Trends and either high Biological Vulnerability or high Action Needs (Gotthardt et al. 2012). All Orange V species scored low on Biological Vulnerability and high on Action Needs. Two of the highest ranked species according to NatureServe, *Myotis lucifugus* (S3, vulnerable) and *Ochotona collaris* (S3S4, vulnerable/apparently secure), were in this category.

Fifty percent of species received a rank of S5 (secure) under the NatureServe criteria, which indicates lowest concern (Droghini et al. 2020). We assigned Blue only to *Lepus americanus*, *Myodes rutilus*, and *Tamiasciurus hudsonicus* because they had low Biological Vulnerability and low Action Needs scores.

### ASRS Theme Scores

#### Population and Distribution Trends

The median score for both questions in Trends was 0, indicating unknown trends. All 36 species had unknown population trends, while 33 species had unknown distribution trends. Distributions of *Lepus americanus*, *Marmota monax*, and *Synaptomys borealis* are known or suspected to have expanded in Alaska over the past fifty years (Tape et al. 2016; A. Baltensperger, pers. comm.; L.E. Olson, pers. comm.).

#### Biological Vulnerability

The median score for Biological Vulnerability was −32, out of a possible minimum of −50. When grouped by order, median scores for Eulipotyphla, Rodentia, and Lagomorpha were low (range: −36 to −32). The median score for Chiroptera was −7, which we consider high. Top-ranking species for Biological Vulnerability were *Sorex pribilofensis* (theme score = 14), *S. jacksoni* (8), and *Myotis volans* (3).

Median scores for range size and number of aggregation sites were the lowest possible scores, indicating that most small mammal species are widespread in Alaska (Fig. 2). Variability in scores for these questions was minimal and characterized by the presence of outliers. The median score for population size was −6 (Fig. 2), which is selected if the population size is unknown but suspected to be large (i.e., more than 10,000 individuals; Gotthardt et al. 2012). Median scores for dietary specialization and habitat specialization were 1; because these questions are assessed on 3-point scales, a value of 1 indicates moderate specialization (Fig. 2). No species was assessed to have high dietary specialization and few species were assessed to have high habitat specialization (Fig. 2).

**Figure 2.**
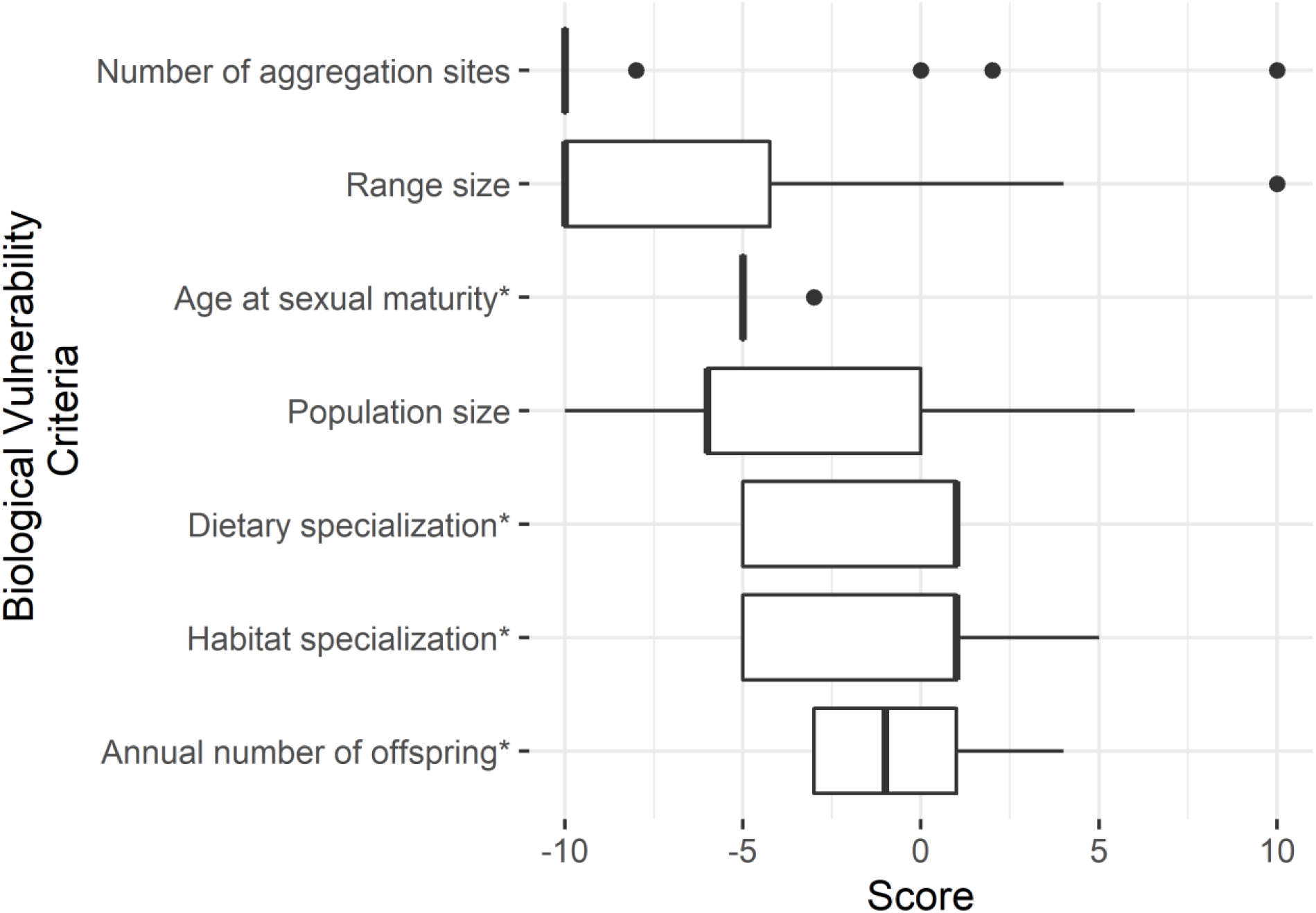
Boxplots of scores in the Biological Vulnerability theme for 36 small mammal species. The bottom and top edges of each box represent the first and third quartiles, respectively. The median is represented by a dark vertical line. Horizontal lines extend no further than 1.5 times the interquartile range. Data points beyond this range are depicted by solid circles. Criteria with an asterisk are evaluated on a scale that ranges from −5 to +5; all other criteria are evaluated on a scale ranging from −10 to +10. Conservation concern increases with numeric value.

#### Management and Conservation Action Needs

No species obtained partial scores for any questions in Action Needs. The median score for Action Needs was 24. When grouped by order, Eulipotyphla had the highest median score (32) while Lagomorpha had the lowest (4). Two species received a score of 40, which is the maximum possible score for Action Needs: *Sorex minutissimus* and *S. navigator*. Most species had high management needs, indicating that they are not subject to direct management actions, and high monitoring needs, indicating that their population trends are not consistently or extensively monitored (Fig. 3). Species with moderate monitoring needs belonged to one of two families: Vespertilionidae or Leporidae. These species are monitored by state agencies, but data are inadequate to detect trends.

**Figure 3.**
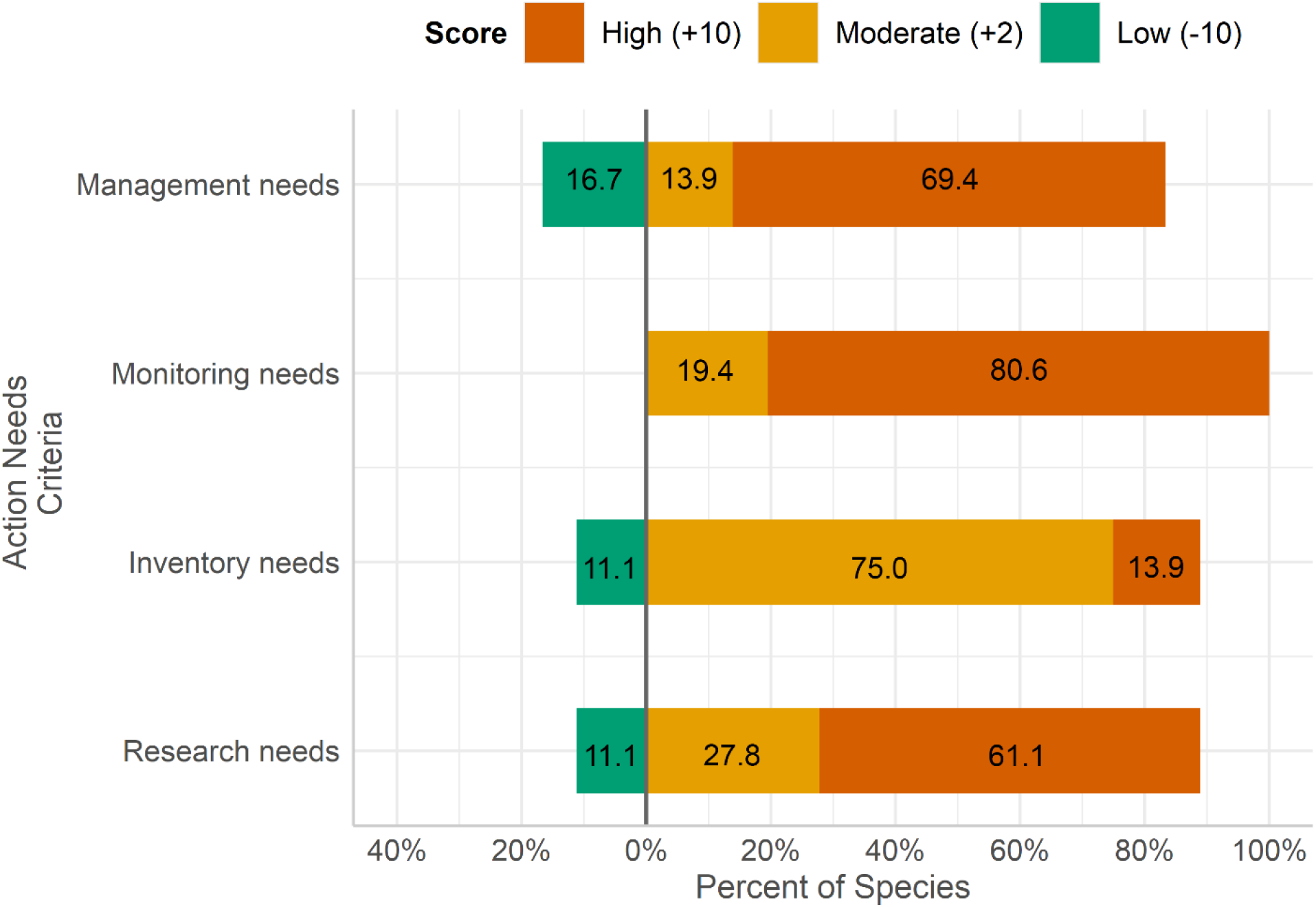
Distribution of ASRS scores for the 4 questions that compose the Action Needs theme. All questions were evaluated on a 3-point scale ranging from −10 to +10; no partial scores were awarded. Conservation concern increases with numeric value.

Nearly all species (*n* = 32) had high or moderate inventory needs; thus, knowledge of range limits and habitat associations remains incomplete (Fig. 3). The species with low inventory needs were *Lepus americanus*, *Myodes rutilus*, *Peromyscus keeni*, and *Tamiasciurus hudsonicus*. These species are widespread, common, and easy to detect or capture in traps.

Twenty-three species (64%) had high research needs, reflecting a lack of information on the factors that limit populations. These species included all species endemic to Alaska, all Eulipotyphla, and all Chiroptera with the exception of *Myotis lucifugus*.

### NatureServe Threats Assessments

Two-thirds of the species we assessed (*n* = 24) received a low threat impact score. Non-native disease (i.e., white-nose syndrome) was listed as a threat for all Chiroptera species, though impact scores varied by species and ranged from very high to medium-low (Droghini et al. 2020). We considered timber harvest to be a medium-low threat for Chiroptera species largely restricted to Southeast Alaska. We considered habitat alteration due to climate change a high-medium threat for talus specialists. *Sorex jacksoni* and *S. pribilofensis* received an impact score of high-low; this score reflects the potentially large, but highly uncertain effects of disturbances on narrowly endemic species.

Experts tended to disagree about the severity or scope of threats, which is reflected for some species as ranges in the overall impact scores (Droghini et al. 2020). Experts also disagreed about the timing of climate change related threats, both within and across species, reflecting uncertainty as to whether effects would be expressed in the short- or long-term (Droghini et al. 2020).

### Recommended Conservation Actions and Research Priorities

We received 23 completed surveys: 5 per species with the exception of *Synaptomys borealis*, for which we were able to identify only 3 experts. The most commonly suggested conservation actions to mitigate threats were to collect more information and to protect known habitat. When asked to allocate US $10 million to different research topics, experts considered monitoring of population trends, research on genetic diversity and adaptive capacity, and habitat modeling important for all species, with $2 to $3 million devoted to each topic (Fig. 4). They considered research on response to climate change important for *Marmota broweri* and *Ochotona collaris*, while research on response to human development and deforestation was important for *Glaucomys sabrinus*, *Myotis lucifugus*, *Synaptomys borealis*. Research on introduced species and on diseases or parasites was judged to warrant comparably little funding (Fig. 4).

**Figure 4.**
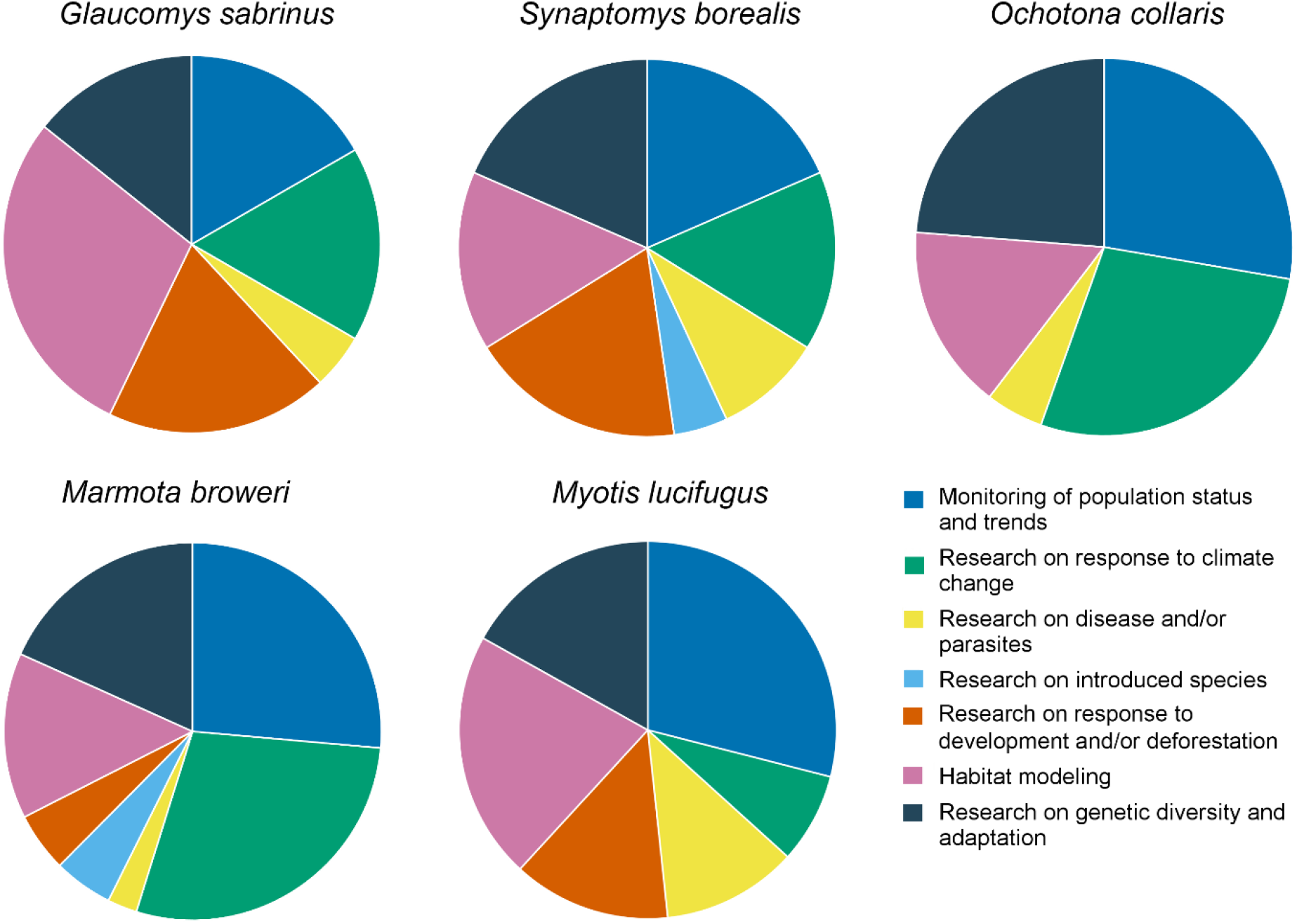
Responses to survey question asking respondents to allocate money to different research topics if given a total budget of $10 million per species. Pie slices represent the proportion of the budget agreed upon by ≥50% of respondents. When there was no consensus among respondents, the median value was used.

## DISCUSSION

The Alaska Species Ranking System (ASRS) explicitly identifies key information needs by assessing the strength of conservation actions around inventory, monitoring, and research. More than 2/3 of small mammal species in Alaska are of high conservation concern in the ASRS, a result driven largely by the lack of information about species’ population trends, distributional limits, and population ecology. The prevalence of data deficiencies for small mammals is not unique to Alaska: relative to other mammal groups, a large proportion of small mammal species are listed as data-deficient by the IUCN but are likely threatened (Jetz & Freckleton 2015).

Species of the orders Chiroptera and Eulipotyphla and species endemic to Alaska were of particularly high conservation concern. Chiroptera species have low reproductive rates and specific habitat requirements for roosting and hibernating; several Chiroptera species also have narrow dietary niches (Safi and Kerth 2004; Boyles and Storm 2007). These traits may contribute to increased extirpation risk (Safi and Kerth 2004; Boyles and Storm 2007). In fact, Chiroptera has experienced a high number of recent extinctions relative to other orders (Ceballos & Brown 1995). All Eulipotyphla species in Alaska received very high Action Need scores in the ASRS; the recent discovery of a species new to Alaska (Dokuchaev 1997; now recognized as *Sorex minutissimus*) and important taxonomic revisions (Hope et al. 2012; Woodman 2018) provide further evidence of high information needs for Eulipotyphla species in Alaska. Data deficiencies in Eulipotyphla also exist at a global scale, despite high levels of diversity and extinction relative to other mammalian orders (Jetz and Freckleton 2015; Verde Arregoitia 2016). Two Eulipotyphla species, *Sorex jacksoni* and *S. pribilofensis*, were the highest-ranked species in both the ASRS and NatureServe. Their ranges are restricted to single islands in the Bering Sea; narrowly endemic species have greater risk of extinction due to small population sizes, small range sizes, and demographic stochasticity (Hartley and Kunin 2003; Cardillo et al. 2008).

In general, the ASRS and NatureServe ranking systems agreed on the rankings of species in the highest and lowest risk categories. However, most species that were of intermediate concern by the ASRS were ranked of lowest conservation concern by NatureServe. Both ranking systems recognize these species’ low biological vulnerabilities: these species are relatively widespread, presumed common, and have life history traits and ecological preferences that correlate with low extirpation risk. The divergence in conservation ranks largely reflects the importance that the ASRS ascribes towards information needs and data deficiencies; in the NatureServe ranking system, data deficiencies do not weight the score towards greater conservation concern.

### Threats to Small Mammals in Alaska

We assessed most small mammal species as having low threat impact scores. Talus specialists and Chiroptera species severely affected by white-nose syndrome received the highest impact scores. In the eastern U.S., populations of *Myotis lucifugus* and *M. septentrionalis* (a close relative of *M. evotis*) declined by over 80% after being infected by *Pseudogymnoascus destructans*, the fungus that causes white-nose syndrome (Langwig et al. 2015). The disease was detected in the western U.S. for the first time in 2016; the taxonomic experts we surveyed expressed uncertainty about the timing of white-nose syndrome (i.e., when it would arrive in Alaska), but predicted strong negative effects to *M. lucifugus*. Based on our literature review, we do not expect other Chiroptera species in Alaska to experience similar population declines from white-nose syndrome.

Talus specialists such as *Marmota broweri* and *Ochotona collaris* occupy habitats that are considered vulnerable to climate change; resulting changes in temperature, snow conditions, and vegetation are expected to affect several aspects of these species’ biology and ecology, including their distribution, thermoregulation, diet, and dispersal (COSEWIC 2011; Hope et al. 2015; Berteaux et al. 2017). At the same time, experts in our survey expressed high uncertainty about the severity, scope, and timing of climate-related threats. It may be possible for talus specialists to adapt and persist by following the movement of alpine plant communities to higher elevations or areas of glacial melt; this spatial shift has been observed in talus specialists in the contiguous United States (Beever et al. 2011). Tundra-adapted species (e.g., *Dicrostonyx groenlandicus*, *Microtus miurus*) may also be threatened by climate change (Lanier et al. 2015; Colella et al. 2020). Unlike talus specialists, which have restricted distributions, tundra-adapted species in Alaska are widespread and often occupy a range of habitats within the broader tundra ecosystem. Moreover, distribution models for these species disagree about the magnitude and direction of climate change effects (Baltensperger and Huettmann 2015; Hope et al. 2015). Thus, for tundra-adapted species, we assumed that the geographic scope of habitat loss due to climate change would affect no more than 30% of the population, and, where habitat loss occurred, it would result in no more than a 30% decline in population. Assumed reductions resulted in a low impact score under the NatureServe methodology (Master et al. 2012). If we were to increase the geographic scope or severity of these threats, the status of these species would increase from S5 (secure) to S4 (apparently secure) in the NatureServe ranking system. ASRS ranks would be unaffected because the ASRS does not include criteria related to threats.

### Recommendations for Conservation Actions and Research Priorities

Effective conservation and management requires accurate knowledge of species’ biology, ecology, and taxonomy (Entwistle & Stephenson 2000). For most small mammal species in Alaska, our understanding of these aspects is severely limited. Indeed, the experts we surveyed identified a need to collect more information for 4 of the 5 species they evaluated. Incomplete knowledge of species’ biology and ecology may lead to incorrect assessments of extirpation risk, and it limits our ability to predict and mitigate the effects of threats. For example, predicting responses to climate change, which experts identified as a research priority, requires a comprehensive understanding of the species of interest, including their ecological requirements, dispersal potential, genetic variability, and phenotypic plasticity (COSEWIC 2011; Colella et al. 2020). Experts selected many of these topics as research priorities.

We identified a vital need to monitor population and distribution trends, which were unknown or uncertain for nearly all species that we assessed. Most small mammals in Alaska are not monitored annually by government agencies. Consequently, the monitoring that is conducted is typically highly localized or only supported for a few years. While preferable to the absence of any monitoring effort, sporadic and isolated monitoring efforts cannot provide robust data on statewide population trends, which require long-term and extensive investments. Although funding for small mammal research is limited, we believe there is considerable potential to develop research programs that address data deficiencies while benefiting existing priority species. Documenting changes in the abundance of small mammals provides valuable insights on the transmission of human diseases and on the ecology of threatened and harvested species such as carnivores, raptors, and waterfowl (e.g., Bêty et al. 2002; Ecke et al. 2017; Schmidt et al. 2018). The value of monitoring small mammal species clearly extends beyond the target species, though this fact is not often recognized by funding agencies or the public.

Small mammals play important ecological roles as herbivores, seed dispersers, and prey. In Alaska, the paucity of data on population size, distribution trends, and basic ecology hinders our ability to assess the health of small mammal populations, including endemic species. Addressing existing data gaps will enable more robust assessments of conservation status for small mammal species and is critical given the rapid pace of climate change and related ecosystems effects.

## ACKNOWLEDGMENTS

We would like to thank the scientists that reviewed the ASRS assessments: A. Baltensperger, K. Blejwas, C. Brandt, A. Gunderson, A. Hope, H. Lanier, L. Olson, and J. Reimer. The following experts participated in the survey: V. Bakker, A. Baltensperger, A. Bidlack, K. Blejwas, D. Causey, E. Flaherty, J. Hagelin, H. Lanier, T. Lee, S. Morrison, T. Mullet, L. Olson, V. Patil, S. Pyare, J. Reimer, K. Rubin, P. Schuette, R. Shively, and W. Smith. T.L. Fields, T.A. Gotthardt, and K.M. Walton made important contributions to earlier versions of the ASRS. M.L. Carlson and T.W. Nawrocki provided valuable feedback on the manuscript. The Alaska Department of Fish and Game and the University of Alaska Anchorage provided financial support for this project.

